# Quantitative profiling of axonal guidance proteins during the differentiation of human neurospheres

**DOI:** 10.1101/2021.02.21.432144

**Authors:** Livia Goto-Silva, Michele Martins, Jimmy Rodriguez Murillo, Leticia Rocha, Gabriela Vitória, Júlia T. Oliveira, Juliana M. Nascimento, Erick Correia Loiola, Fabio C. S. Nogueira, Gilberto B. Domont, Marília Zaluar P. Guimarães, Fernanda Tovar-Moll, Steven Kastrup Rehen, Magno Junqueira

## Abstract

Axon guidance is required for the establishment of brain circuits. Whether much of the molecular basis of axon guidance is known from animal models, the molecular machinery coordinating axon growth and pathfinding in humans remains to be elucidated. The use of induced pluripotent stem cells (iPSC) from human donors has revolutionized *in vitro* studies of the human brain. iPSC can be differentiated into neuronal stem cells which can be used to generate neural tissue-like cultures, known as neurospheres, that reproduce, in many aspects, the cell types and molecules present in the brain. Here, we analyzed quantitative changes in the proteome of neurospheres during differentiation. Relative quantification was performed at early time points during differentiation using iTRAQ-based labeling and LC-MS/MS analysis. We identified 6,438 proteins, from which 433 were downregulated and 479 were upregulated during differentiation. We show that human neurospheres have a molecular profile that correlates to the fetal brain. During differentiation, upregulated pathways are related to neuronal development and differentiation, cell adhesion, and axonal guidance whereas cell proliferation pathways were downregulated. We developed a functional assay to check for neurite outgrowth in neurospheres and confirmed that neurite outgrowth potential is increased after 10 days of differentiation and is enhanced by increasing cyclic AMP levels. The proteins identified here represent a resource to monitor neurosphere differentiation and coupled to the neurite outgrowth assay can be used to functionally explore neurological disorders using human neurospheres as a model.

## 1. Introduction

Axon guidance is one of the first steps in the formation of brain circuits [1,2]. This process is chemically regulated by attractive and repulsive cues, which are secreted by target organs and triggers signaling at the neuronal growth cone with extensive cytoskeletal remodeling.

The *in vitro* study of human brain tissue using cells containing the same genome as human donors was made possible since the discovery that somatic cells can be induced into a pluripotency state. The induced expression of four transcription factors, Klf4, Oct3 / 4, Sox2 and cMyc, also known as Yamanaka factors, in somatic cells, can generate induced pluripotent stem cells (iPSC), which present features of human embryonic cells, such as the ability to differentiate into cells of the three germ layers [3–5]. This technique has opened new perspectives for the exploration of translational therapies, in which cultures of cells from patients can be used for drug screening and personalized medicine. Especially for the study of the brain, this advance is significant, given the obvious difficulty of obtaining live human central nervous system (CNS) samples.

Renewable populations of neural stem cells (NSC) can be obtained from iPSC maintained in chemically defined culture media. These cells can be differentiated to neural cell types, including neurons, astrocytes, and oligodendrocytes [6–8] and have been proven useful to study several neurological conditions, such as Amyotrophic lateral sclerosis, schizophrenia, Rett syndrome, and Alzheimer’s disease [9–13]. Despite iPSC models contribution to the study of human diseases, most of the data generated were based on cells growing in two-dimensional (2D) cultures, where cell-cell contacts are limited and, hence, lack spatial tridimensional (3D) information which is important for tissue development.

Cultivation of NSCs into free-floating spheric clusters, which are induced for neural differentiation, and therefore termed neurospheres, leads to a more representative model of the cell-cell interactions and the microenvironment of the brain tissue [14]. Neurospheres have been used to study the developing brain [15,16], and recent advancements also made it possible to address drug neurotoxicity and neural activity in this culture system [17,18]. Despite the relevance of neurospheres for disease modeling, a comprehensive view of proteins and pathways present in these cells is lacking. Mass spectrometry-based quantitative proteomics has emerged as a powerful tool to quantify proteins and identify pathways altered in complex biological systems [19]. Therefore, in this work, we applied isobaric tags (iTRAQ) to quantify proteins during the early differentiation of human neurospheres. We show that whereas proliferation pathways decrease with time of differentiation, axonal guidance pathways are increased, which is confirmed by a neurosphere neurite outgrowth assay. We propose that the data obtained with this proteomic pipeline together combined with an axon outgrowth assay are useful tools to address drug discovery for human neurological pathologies modeled by iPSC technology.

## 2. Material and methods

### 2.1. Cell culture

Human iPSCs were obtained from the Coriell Institute for Medical Research repository (GM23279A) or produced in house with CytoTune^™^-iPS 2.0 Sendai Reprogramming Kit (Invitrogen) from skin fibroblasts [20] or urine epithelial cells [21]. Human iPSCs were maintained in Stem Flex media (Thermo Fisher Scientific), on Matrigel (BD Biosciences) coated dishes, at 37 °C in humidified air with 5% CO2. Cultivation media was changed daily to avoid differentiation of cultures. The colonies were manually split upon reaching 70–80% of confluence. Differentiation to NSCs was induced by cultivation in PSC neural Induction Medium, containing Neurobasal medium and neural induction supplement (Thermo Fisher Scientific), according to manufacturer’s protocol. Media was changed every other day until day 7, during which NSCs were split and expanded on neural expansion medium (1:1, Advanced DMEM/F12 and Neurobasal medium) with neural induction supplement (Thermo Fisher Scientific).

Neurospheres for proteomic analysis were prepared in suspension as follows: upon reaching 90% of confluence, NSCs were detached from the plate using accutase (Merck Millipore), centrifuged at 300 xg for 5 minutes, and resuspended in neural induction medium with 1:1 DMEM / F12 (Life Technologies) and Neurobasal medium (Thermo Fisher Scientific) supplemented with 1x N2 (Invitrogen) and 1x B27 (Thermo Fisher Scientific). 3 x 10^6^ NSCs were plated in each well of a 6 well-plate and cultured on an orbital shaker at 90 rpm for 3 or 10 days at 37 °C in humidified air with 5% CO2. The medium was changed each 3-4 days [22].

### 2.2. Protein extraction for proteomics

Neurospheres were washed twice with cold PBS and centrifuged at 300 xg for 5 minutes. Residual liquid was removed before storage at −80 °C until further processed. Neurospheres formed from 3 x 10^6^ cells were lysed with buffer containing 7 M urea, 2 M thiourea, 50 mM HEPES pH 8, 75 mM NaCl, 1 mM EDTA, 1 mM phenylmethylsulfonyl fluoride, and protease/phosphatase inhibitor cocktails (Roche). The lysates were sonicated and centrifuged at 10,000 xg for 10 min at 4 °C. Total protein content in the supernatant was quantified with Qubit^®^ fluorometric assay (Invitrogen) following the manufacturer’s instructions. Two independent biological replicates of each condition were used in the experiments.

### 2.3. Protein digestion and iTRAQ labeling

Proteins (100 μg each condition) were incubated with dithiothreitol at 10 mM final concentration for 1 h at 30 °C; afterward, iodoacetamide (final concentration of 40 mM) was added and the samples were incubated for 30 min in the dark at room temperature. Samples were diluted 10-fold with 50 mM HEPES pH 8 and incubated with sequence-grade modified Trypsin (Promega) at 1/50 trypsin/protein ratio for 16 h at 37 °C. The resultant peptides were further labeled using isobaric tags for relative and absolute quantification (iTRAQ) 4-plex commercial reagent kit (AB Sciex) according to the manufacturer’s instructions. Peptides from two biological replicates of each condition were labeled as follows: 114 and 116 isobaric tags for 3-day old neurospheres; 115 and 117 for 10-day old neurospheres. Labeled peptides were mixed in a unique tube and desalted with a C-18 macro spin column (Harvard Apparatus) and then dried in a vacuum centrifuge.

### 2.4. Hydrophilic interaction chromatography fractionation and nano-LC–MS/MS analysis

Dried peptides were pre-fractionated offline using hydrophilic interaction chromatography (HILIC). Peptides were redissolved in buffer A (90% acetonitrile, 0.1% trifluoroacetic acid) and then fractionated using a UPLC System (Shimadzu) connected to a HILIC-TSKGel Amide-80 column (5 cm × 2 mm i.d. × 5 μm) (Supelco). The separation was achieved with a linear gradient of 0 to 30% of buffer B (0.1% trifluoroacetic acid) in 45 min at a constant flow rate of 200 μl/min. A total of 26 fractions were collected (those with low absorbance intensity were pooled, resulting in 9 fractions) and dried in a speed vacuum concentrator. Each fraction or pool of fractions was analyzed in three technical replicates in an Easy-nLC 1000 nano-LC system (Thermo Scientific) coupled to a Q-Exactive Plus mass spectrometer (Thermo Scientific). Fractions were dissolved in 20 μl of 0.1% formic acid, loaded onto a trap column (ReprosilPur C18, 2 cm × 150 μm i. d. × 5 μm) with a flow rate of 5 μl/min and separated on the analytical column (ReprosilPur C18, 30 cm × 75 μm i.d. × 1.7 μm) with a constant flow rate of 300 nL/min and a linear gradient of 5–40% of B (95% acetonitrile, 0.1% formic acid) in 120 min. The mass spectrometer was set as follows: electrospray was used at 2.0 kV and 200°C, data acquired in dependent analysis mode with dynamic exclusion of 30 s, full-scan MS spectra (*m/z* 375–1800) with a resolution of 70,000 (*m/z* 200), followed by fragmentation of 12 most intense ions with high energy collisional dissociation, normalized collision energy of 30 and resolution of 17,000 (m/z 200) in MS/MS scans. Species with a charge of +1 or greater than +4 were excluded from MS/MS analysis.

### 2.5. Data analysis

Raw data were processed using Proteome Discoverer 2.4 Software (Thermo Scientific). Peptide identification was performed with the Sequest HT algorithm against the Human proteome database provided by Uniprot (http://www.uniprot.org/downloaded in 2020 26-04-2016). The searches were run with peptide mass tolerance of 10 ppm, MS/MS tolerance of 0.1 Da, tryptic cleavage specificity, 2 maximum missed cleavage sites, fixed modification of carbamidomethyl (Cys), and variable modification of iTRAQ 4-plex (Tyr, Lys, and N-terminus), and oxidation (Met), false discovery rates (FDR) were obtained using Target Decoy PSM Validator node selecting identifications with a *q*-value equal or to less than 0.01.

Data was quantified in software Perseus [23]. Data from all technical and biological replicates were transformed to log2 and normalized by subtracting the median. Technical replicates were averaged, and proteins were considered differentially expressed if the 3-day and 10-day ratios (10D / 3D) were ≥1.5 or ≤0.67 thresholds if the iTRAQ ratios (115/114 and 117/116) with 95% confidence values (significant p<0.05) from student’s t-test. The Biological processes and pathways enrichment for regulated proteins was performed using the online tool metascape (https://doi.org/10.1038/s41467-019-09234-6), available at https://metascape.org/ or the tool Enrichr [24](https://maayanlab.cloud/Enrichr/). Metascape utilizes the well-adopted hypergeometric test58 and Benjamini-Hochberg p-value correction algorithm [25]. All analysis was performed with P-Value Cutoff 0.01 using data from Gene Ontology (GO) [26] and KEGG. The PSM values (Log2 transformed) of neurospheres were compared with data of cerebral organoids proteomics [27], and with proteomics data of fetal and adult human brain cortex [28], using Pearson correlation.

### 2.6. Neurite outgrowth assay

Neurospheres were formed by plating 9 x 10^3^ NSCs in 150 mL of round bottom Ultra-low attachment 96 well plates (Corning). After plating, cells were centrifuged 300 x*g* for 3 minutes to assure sedimentation. To access neurite outgrowth, one neurosphere was plated per well of 96 well plates (PerkinElmer) previously coated with 100 μg/ml polyornithine and 20 μg/ml laminin at days 3 and 10. Neurites were allowed to grow for 48 hours. 10-day old neurospheres were treated with 10 μM forskolin (Alomone Labs) 24 hours after plating and analyzed 24 h after treatment. Neurospheres were fixed with 4% paraformaldehyde solution for 20 minutes.

Immunohistochemistry was performed as follows: fixed neurospheres were washed with PBS, permeabilized with 0.3% Triton X-100 in PBS for 15 min. and incubated in a blocking solution consisting of 3% goat serum in PBS. Cells were permeabilized with 0.3% Triton for 30 min followed by the primary antibody (Tuj 1, M015013, Neuromics) solution (1:4000) overnight at 4°C. Then, neurospheres were washed 3 times with PBS and incubated with secondary antibody goat anti-mouse Alexa Fluor 488 (Thermo Fisher Scientific) diluted at 1:1000 for 45 minutes. Nuclei were counterstained with 0.5 μg/mL 40-6-diamino-2-phenylindole for 5 minutes, glycerol was added to plates, and images were acquired in the Operetta High Content Analysis System (PerkinElmer) using 2x or 10x objectives. Neurite growth, analyzed by the total length and number of neurites parameters, was quantified using the high-content image software Harmony 5.1 (PerkinElmer) and the neuronal morphology analysis tool of WIS-NeuroMath software, respectively. For quantification by Neuromath, a circular mask was created at 4 times the diameter of the neurosphere and the number of neurites crossing this mask was quantified. Differences were analyzed by non-parametric t-test, followed by the Mann Whitney test, using GraphPad Prism software 7.0. A confidence interval of 95% was accepted as significant (*p*<0.05).

## 3. Results and discussion

### 3.1. Neurosphere proteins overlap fetal brain proteins

The proteome of human neurospheres was quantitatively analyzed at two-time points after the formation of neurospheres. After neurosphere formation, the first time point of manipulation for media change is day 3. This time point was chosen to start the study of neurosphere differentiation. A later time point, 10 days after differentiation, was previously reported to derive neurospheres that, after plating, can migrate and project axons radially, eventually forming functional neural networks [29]. Therefore, 10 days after plating was chosen as the second time point to study neurosphere differentiation. Neurospheres were analyzed after 3 or 10 days in suspension culture with differentiation media in 2 biological replicates. Proteins were trypsin digested and peptides were labeled with ITRAQ and mixed in a 1: 1: 1: 1 ratio. Peptides were fractionated off-line with HILIC and fractions were analyzed by LC-MS / MS (Supp. Fig. 1). Datasets were submitted to statistical analysis based on the values of the relations of channels 115/114 and 117/116, analysis of enrichment of GO terms, and network of the interaction of key proteins (Fig. 1A). Typical morphology of neurospheres of 3 and 10 days is represented in figure 1B.

**Fig.1:**
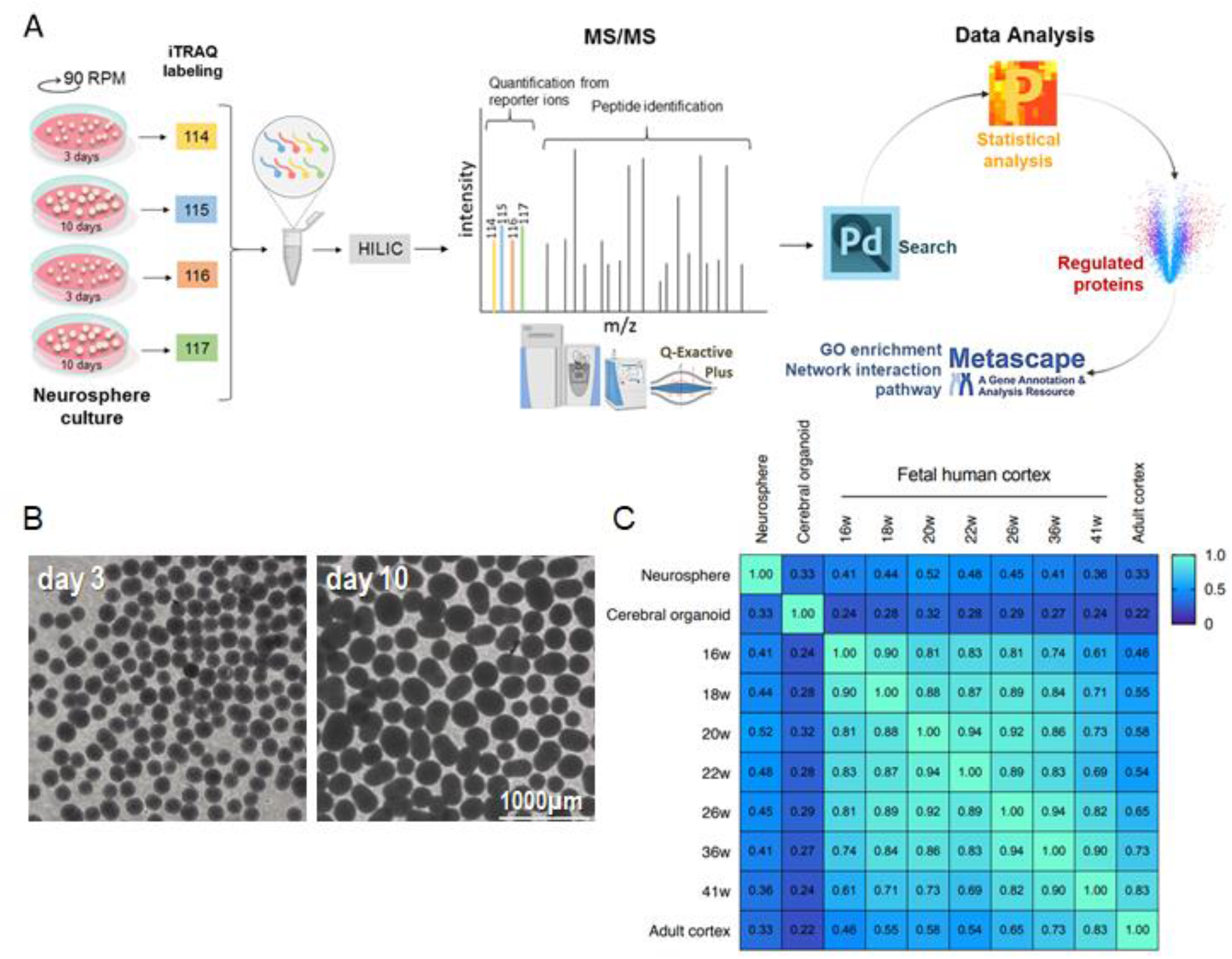
Identification of proteins in neurospheres shows overlap with fetal brain proteins. A) Quantification pipeline. Neurospheres in rotational cultures for 3 or 10 days were solubilized, proteins were quantified and digested, and peptides were labeled using an iTRAQ-4-plex reagent. Samples were pulled together and fractionated in HILIC and fractions were analyzed in a Q-Exactive orbitrap. Peptides were quantified using the reporter ion signal and identified based on their fragmentation spectra. Data were analyzed in Proteome Discoverer 2.4 and statistics were performed in Perseus 1.6.12. GO enrichment was carried out in Metascape online tool. B. Neurospheres cultivated for 3 days and 10 days. C. Pearson correlation of neurospheres proteome to cerebral organoids and 16-41 weeks fetal brain and adult cortex proteome.

The proteome of the pool containing neurospheres of 3 and 10 days (Supp. dataset) was correlated with protein expression data from fetal brains and human brain organoids (Fig. 1C). We could see that the highest correlation is at embryonic week 20 (r=0.52), with the lowest correlation to the adult cortex (0.34). It shows that the molecular profile of neurospheres is more similar to the human cortex at embryonic week 20. A similar correlation of neurospheres and organoids (range r=0.29-0.37, p< 1.6×10-57) is observed with organoids and the fetal human cortex at week 20 of development (range r=0.27 −0.36, p< 1.4×10-25) (Nascimento, 2019). Neurospheres form fewer complex structures than brain organoids, which develop tissue layers similar to those found in the developing cortex. Still, proteins expressed in neurospheres can correlate to the fetal cortex at similar correlation coefficients (ranging from 0.36-0.52, p< 3.6×10-65, depending on the gestational week).

Correlation of the neurosphere protein profile to the protein profile of the fetal cortex validates the use of neurospheres to study the human brain. It is important to highlight that the preparation of neurospheres can be as fast as 3 days, whereas brain organoids are generally used from 30 days in culture. This makes neurospheres attractive as platforms for drugs and disease screenings.

### 3.2. During differentiation, neuronal proteins are increased and cell cycle proteins are downregulated

Proteomic analysis identified 6,438 groups of proteins considering a 1% FDR. Proteins with abundance changes considering a p-value <0.05 are depicted in a volcano plot (Fig. 2A; Supp. dataset). From 3 to 10 days of neurosphere differentiation, a total of 433 proteins were downregulated and 479 were upregulated considering a threshold of log2 fold change 0.5. GO enrichment for biological processes revealed that proteins related to the cell cycle are downregulated (Fig. 2B), whereas proteins that are related to neuronal maturation are upregulated (Fig. 2C).

**Fig.2:**
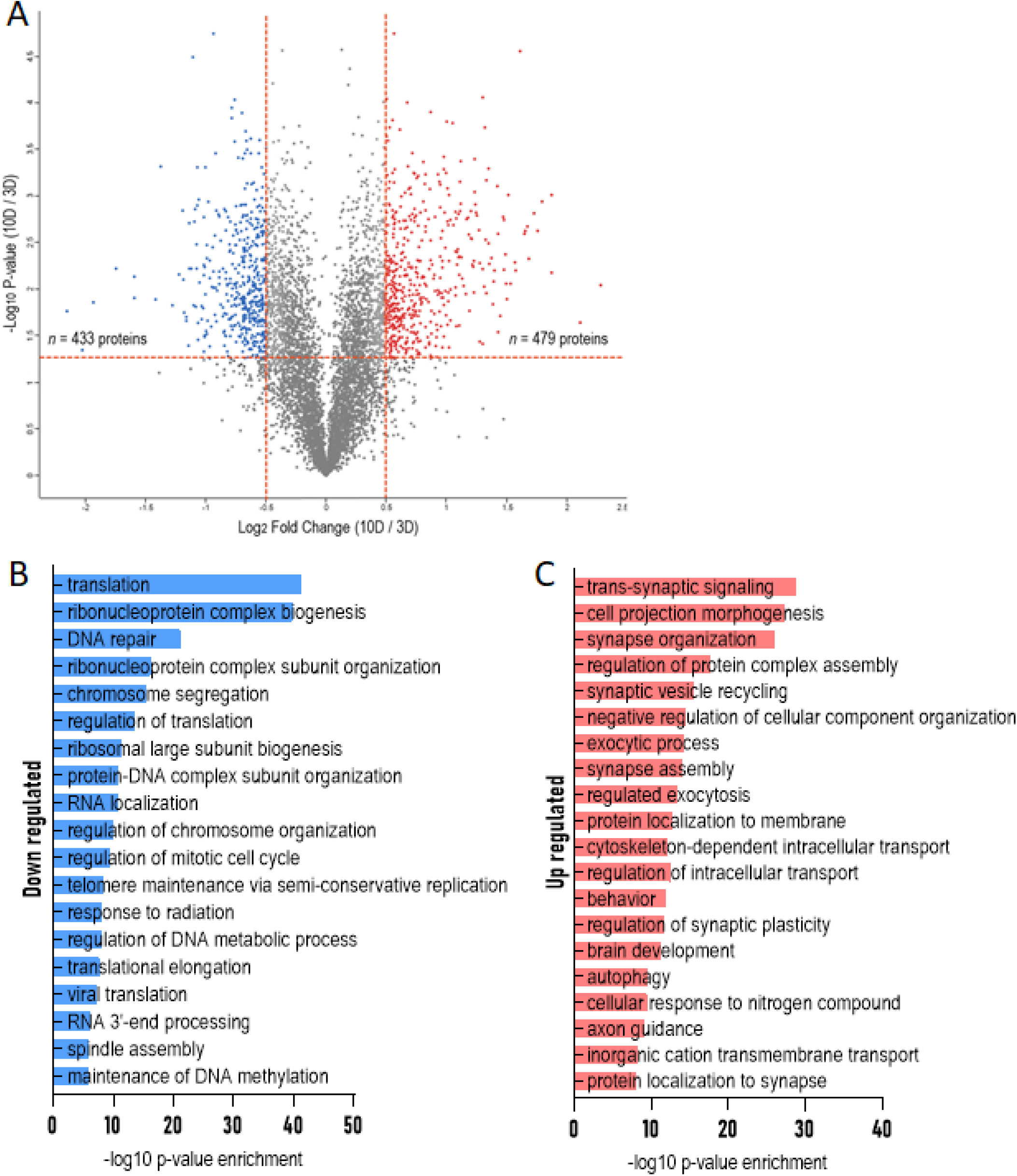
Proteins differentially expressed and significantly enriched biological processes by GO terms between 3 days and 10 days of differentiation. A: Volcano plot displaying the protein abundance changes. The vertical axis corresponds to the log p-value negative of the t-test and the horizontal axis displays the log2 fold-change value. Proteins were considered upregulated and downregulated if fold-change values were ≥1.5 or ≤0.67 at the threshold p-value <0.05. B: Overview of biological processes enrichment [−log10 (P-value)] of downregulated genes in a bar chart. C: Overview of biological processes enrichment [−log10 (P-value)] of upregulated genes in a bar chart.

### 3.3. During differentiation axon guidance proteins are upregulated and neurite outgrowth is increased

We showed that the dataset of proteins upregulated at 10-day old neurospheres is enriched for axon guidance proteins (Fig. 2). Further examining the complete dataset, we observed that out of 96 proteins in the axon guidance pathway (KEGG database), 42 were identified in our dataset, from which 16 were upregulated (Fig. 3). There were no down-regulated proteins from the dataset in the axon guidance pathway. Regulated proteins in the axon guidance pathway are part of the Netrin, Slit, Semaphorins, Netrin-G2, and cell adhesion molecules (CAMs) signaling.

**Fig.3:**
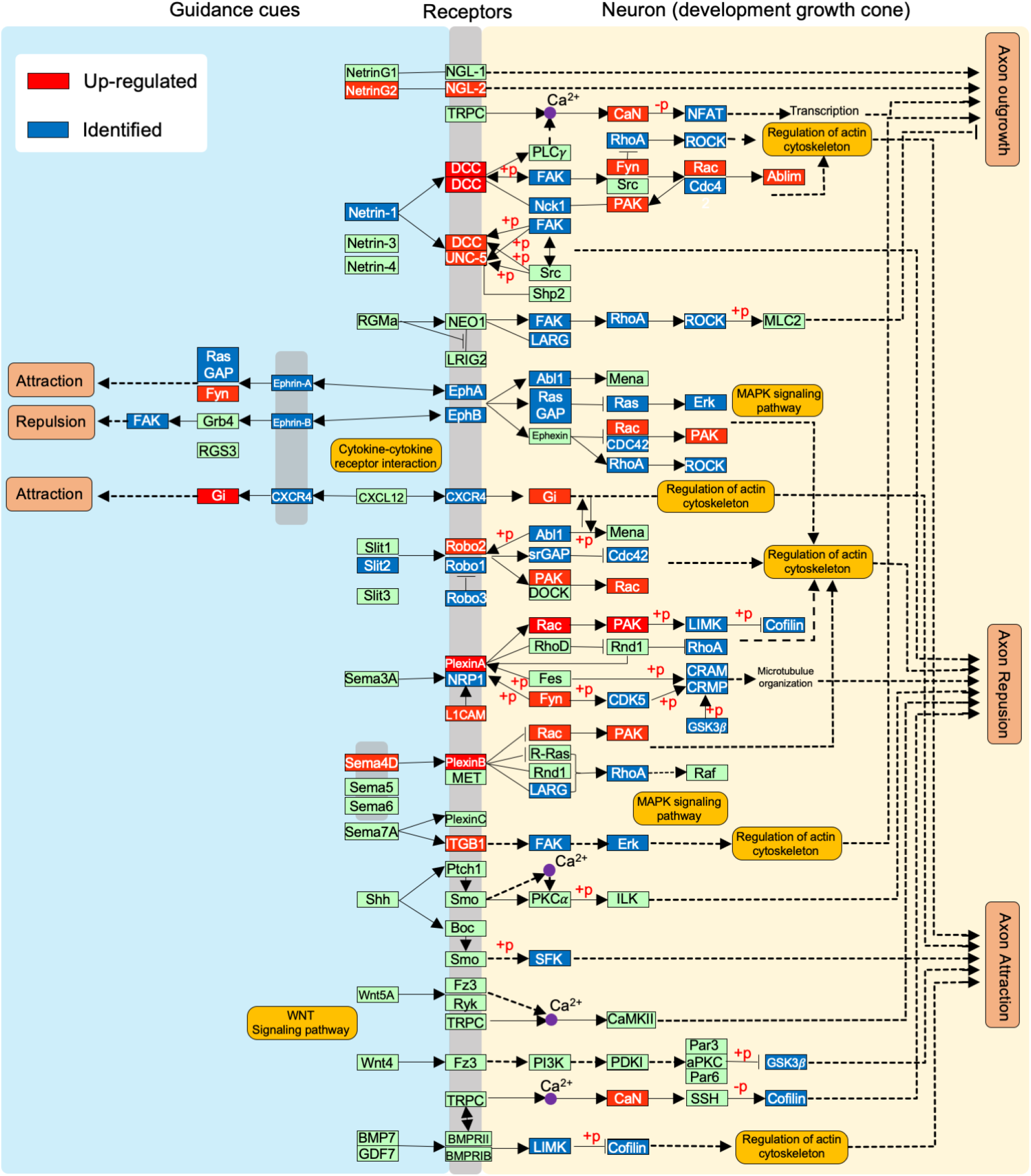
Axon guidance pathway. Proteins related to this pathway detected in our data set are highlighted as identified only (blue) and upregulated (red). Blocks in green represent proteins that were not detected in the dataset. Continuous arrows represent direct molecular interaction or relation. Dotted arrows represent indirect effect or state change. +p: phosphorylation. −p: unphosphorylated.

Netrin 1 was the first axon guidance cue identified in the spinal cord [30]. Netrin produced by the floor plate attracts commissural axons by forming a gradient, with the highest concentration next to the floor plate and also in the central canal [1]. Netrin signaling may promote axon outgrowth or repulsion, depending on whether the two Netrin receptors, UNC5A and DCC are expressed in cells. Binding to DCC, isolated, promotes axon outgrowth, but the association of UNC5A and DCC triggers signaling for repulsion [30,31]. During brain development, repulsion is necessary to guide axons to their proper targets. For example, Slits are secreted by the midline, a lining of glial cells separating the two brain hemispheres. Neurons expressing the receptor ROBO turn away from the midline whereas neurons expressing low levels of ROBO can cross the midline. After crossing the midline neurons increase ROBO expression, this prevents the returning of axons to their hemisphere of origin (Kidd, 1999; Kidd, 1998). Semaphorins also act as chemorepellent molecules when binding to plexin receptors, in association with neural cell adhesion molecule L1 (L1CAM1) [2,32]. Increased expression of UNC5A, DCC, ROBO2, and Plexins suggests that during differentiation, neurospheres acquire the molecular machinery to respond to axon guidance cues.

We set up a neurosphere neurite outgrowth assay to validate whether the increased expression of axonal guidance proteins functionally corresponds to increased neurite migration (Fig. 4A). Neurospheres were formed by plating 9 x 10^3^ NSCs in round bottom Ultra-low attachment 96 well plates to generate spheres of equal size (Fig. 4B). Then, one neurosphere was plated per well of a 96 well plate at 3 and 10 days. Neurite outgrowth was analyzed 24 hours after plating. We compared neurosphere migration at 3 and 10-day after the start of differentiation (Fig. 4C) and observed that 36.6% of the 3-day old neurospheres displayed neurite growth compared with 10-day old neurospheres, from which 85% displayed neurite outgrowth (Fig. 4D). Also, 3-day old neurospheres showed significantly fewer neurite projections than 10-day old neurospheres, as quantified by the total length of the neurites per neurospheres (3460 ± 548.8 vs. 5000 ± 665.5 with a P-value 0.0032) (Fig. 4E). These results confirm that 10-day old neurospheres have increased neurite growth potential and suggest that the axonal guidance genes, which have increased expression at day 10 of differentiation are molecular players in this process.

**Fig.4.**
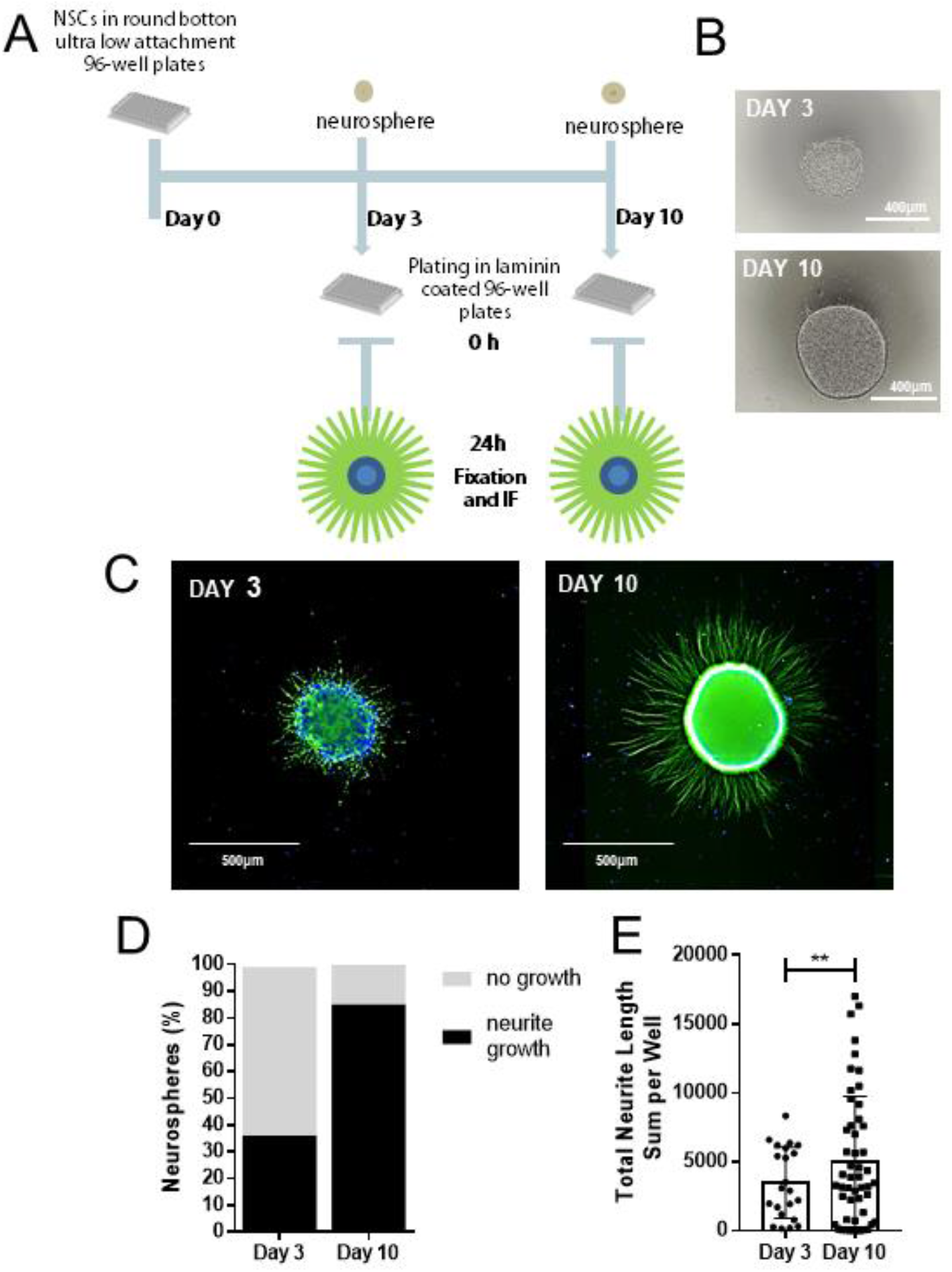
Neurite outgrowth increases during differentiation. A. Neurite outgrowth assay: neurospheres are plated in 96-well plates and neurites are allowed to grow for 24 h. B. 3-days and 10-days neurospheres, 24h after plating before fixation. C. 3-day and 10-day neurospheres, 24 h after plating, immunostained with Tuj-1, nuclei were stained with Dapi. D. Quantification of the percentage of 3- and 10-neurospheres which showed neurite growth. E. Total length of neurites in each neurosphere (**p<0.005). Quantifications are from n=2 and a total of 60 neurospheres.

### 3.4. Regulation of neurite outgrowth in human neurospheres

We showed that neurospheres express netrin receptors, and project neurites radially after plating. Next, we asked whether the neurite outgrowth observed is responsive to regulatory kinases, which have been demonstrated to modulate axon guidance receptors. Appropriate localization of receptors at the growth cone is required for proper response to ligands. It has been described that cAMP and PKA signaling are involved in the transport and localization of netrin receptor DCC at the plasma membrane [33]. An increase in intracellular cAMP, with subsequent activation of PKA, augments the extension of axons from commissural neurons in response to netrin-1 [33]. To verify if an increase in cAMP levels could have a similar effect on human neurospheres, we treated plated neurospheres with forskolin (Fig. 5). Forskolin increases cAMP levels by directly activating adenylate cyclase, which leads to PKA activation. Plated neurospheres were treated for 24 h with 10 μM forskolin, and neurite outgrowth was assayed (Fig. 5A). We observed that PKA activation induced an increase of 138.18 % in neurite growth in comparison to untreated control neurospheres (Control: 11 ± 9.72 and Forskolin-treated: 26.2 ± 11.27, p<0.01) (Fig. 5B). Our data suggest that neuronal guidance proteins detected in the proteomic analysis are functional and responsive to cAMP signaling.

**Fig.5:**
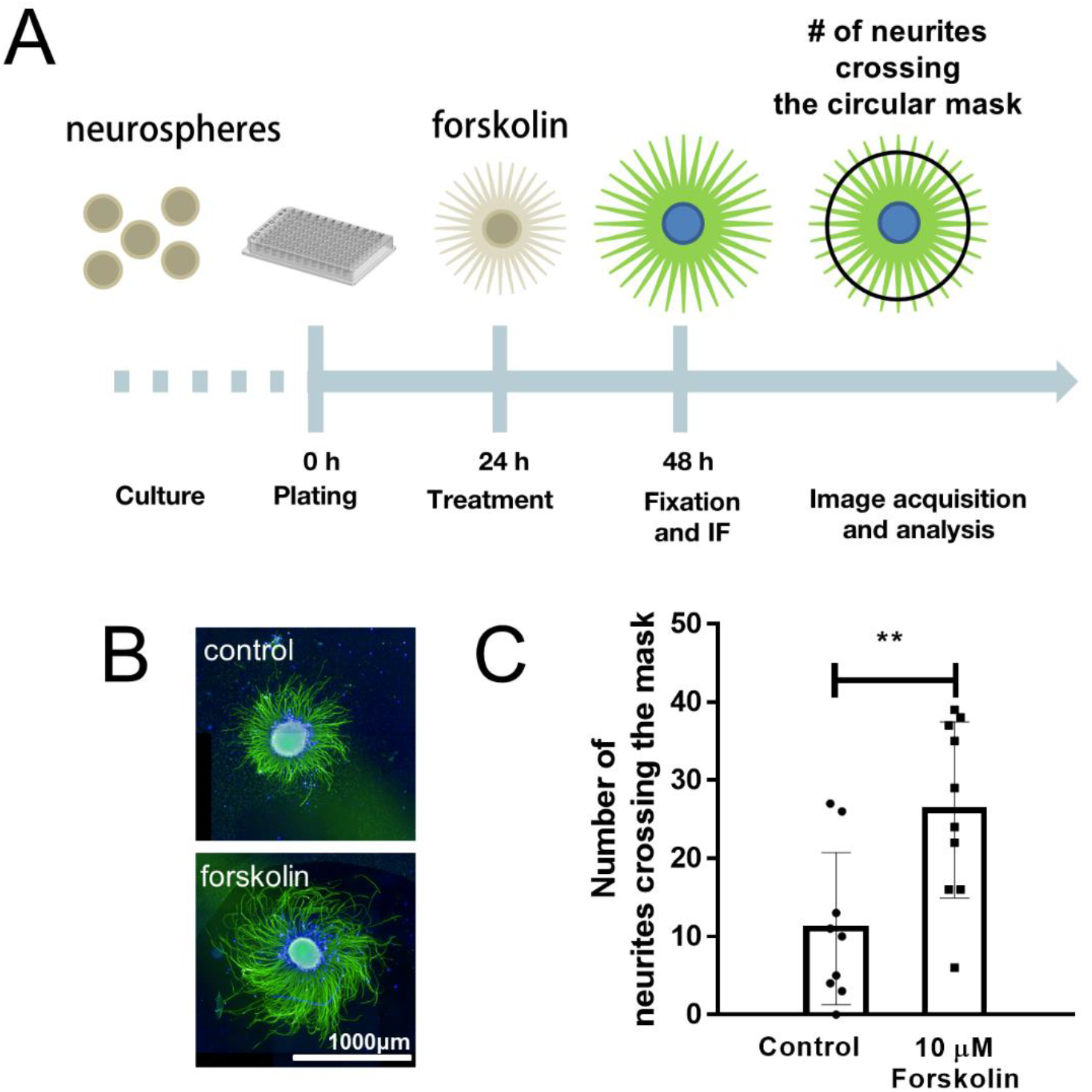
Neurite outgrowth is regulated by cAMP signaling. A. Plated neurospheres were treated with 10 μM forskolin for 24 h. The number of neurites crossing a circular mask around the neurosphere was used for quantification. B. Micrographs of control and 10 μM forskolin-treated neurospheres immunostained for beta III tubulin (green). Nuclei counterstained with DAPI (blue). C. Number of neurites crossing the circle (**p<0.01). Quantifications are from n=1 and a total of 10 neurospheres per group.

### 3.5. Axon guidance regulated genes in human neurospheres are associated with neurological disorders

Several genes from the axon guidance pathway are involved in neurological disorders (reviewed in [34]). To address the future potential to use neurospheres to explore axon guidance proteins in disorders affecting the human brain, we searched the list of proteins identified in the axon guidance pathway in DISGENET and ARCHS^4^ data banks using the Enrichr tool [24]. Table 1 shows axon guidance genes identified in neurospheres with significant overlap to neurodegenerative, neuropsychiatric, and neurodevelopmental disorders. These genes can be divided into membrane receptors, signaling effectors, and kinases.

**Table 1:** Axon guidance genes co-related with neurological disorders. Axon guidance genes identified in our dataset were searched in DISGENET (neurodegenerative and neuropsychiatric disorders) and ARCHS^4^ (neurodevelopmental disorders) using Enrichr tool.

| Neulorogical disorder | Genes |
| --- | --- |
| **Neurodegenerative** |  |
| Alzheimer's Disease | ITGB1; GSK3B; EPHA4; ROCK1; ROCK2; LRRC4; CXCR4; NTN1; RHOA; GNAI1; CDC42; PPP3CA; PPP3R1; CDK5; DPYSL2; CFL2; ABL1; MAPK1; FYN; RAC1; EPHB2; EPHB1; MAPK3; PLXNA3 |
| Parkinson Disease | GSK3B; ROCK2; CXCR4; CDC42; CDK5; RASA1; ABL1; MAPK1; FYN; RAC1; SRGAP3; EPHB2; EPHB1; MAPK3 |
| Huntington Disease | GSK3B; ROCK1; CDK5; DPYSL2; LIMK1; MAPK1; RAC1; EPHB2; EPHB1 |
| **Neuropsychiatric** |  |
| Schizophrenia | GSK3B; DCC; NFATC3; CXCR4; L1CAM; EFNB2; CDC42; PPP3CA; PPP3CB; PPP3R1; CDK5; DPYSL2; CFL1; PLXNA2; MAPK1; FYN; RAC1; SRGAP3; EPHB2; PAK3; EPHB1; PAK2; MAPK3 |
| Major Depressive Disorder | NRP1; GSK3B; CDK5; MAPK1; RAC1; EPHB2; EPHB1; MAPK3; PLXNA3 |
| Bipolar Disorder | GSK3B; PPP3CB; NRAS; DPYSL2; MAPK1; FYN; RAC1; EPHB2; MAPK3 |
| **Neurodevelopmental** |  |
| Agyria-pachygyria type 1 | ROBO2; GSK3B; DPYSL5; DCC; DPYSL2; SRGAP3; PAK3; L1CAM; PAK5; ROBO1; PLXNA3 |
| Corpus callosum dysgenesis X-linked recessive | ROBO2; GSK3B; DPYSL5; DCC; DPYSL2; PLXNA1; PAK3; PAK5; ROBO1; PLXNA3 |
| Macrogyria pseudobulbar palsy and mental retardation | ITGB1; GSK3B; ARHGEF12; DPYSL5; DPYSL2; PLXNA1; PAK3; L1CAM; ROBO1 |
| Cerebellar hypoplasia | GSK3B; DPYSL5; DCC; PLXNA1; SRGAP3; L1CAM; ROBO1; PLXNA3 |
| Familial band heterotopia | GSK3B; ARHGEF12; DPYSL5; DPYSL2; PAK3; L1CAM; ROBO1 |
| Lissencephaly syndrome type 1 | GSK3B; ARHGEF12; DPYSL5; PLXNA1; PAK3; L1CAM; ROBO1 |

L1CAM is highly enriched on the axonal surface of developing and regenerating axons. During brain development, it is critical in multiple processes, including neuronal migration, axonal growth and fasciculation, and synaptogenesis. In the mature brain, L1 CAM plays a role in the dynamics of neuronal structure and function, including synaptic plasticity, and is involved in axonal pathfinding guidance and branching, and myelination. L1CAM is altered in neurodevelopmental disorders and may be regulated upon dysfunction of brain circuitry. A proteomic study of nascent proteins in axons in vitro has demonstrated that L1CAM is translated in axons in response to netrin, semaphorin, and BDNF [35]. This study shows that the fast production of proteins is relevant during axonal outgrowth and could not be estimated by transcriptomic analysis.

It would be promising to address whether L1CAM and receptors for axon guidance cues are regulated or mis localized in models of the diseases listed in Table 1. Also, the signaling cascades triggered by these receptors could be studied at the molecular level in human cells and serve as drug targets for remodeling of the brain circuitry, with possible application in neurodegenerative and neuropsychiatric diseases.

## 4. Conclusions

Here we combine high content data of protein and pathways, and functional assays of processes regulated during neurospheres differentiation. We observed that at early time points of neural induction, several proteins regulating the synaptic formation, matrix attachment, and axon guidance have their expression increased. Data presented here provide an explorer guide to use neurospheres as a comprehensive system to address cell-cell interactions in the context of diseases in which neural circuits are affected. Proteins reported here can also serve as a resource to monitor the effects of drugs in the CNS, thus paving the way for the discovery of mechanisms involved in setting up brain connections.

## Supporting information

proteomic dataset

## Abbreviations

2D: two-dimensional
3D: tridimensional
AMPc: cyclic adenosine monophosphate
CAMs: Cell adhesion molecules
CNS: central nervous system
FDR: false discovery rates
GO: Gene ontology
HILIC: hydrophilic interaction chromatography
iPSC: induced pluripotent stem cells
iTRAQ: Isobaric tag for relative and absolute quantitation
MS: mass spectrometry
NSC: neural stem cells
PBS: phosphate-buffered saline
PKA: Protein kinase A

## Availability of raw files

The mass spectrometry proteomics data have been deposited to the ProteomeXchange Consortium via the PRIDE (https://www.ebi.ac.uk/pride/) partner repository with the dataset identifier PXD024177.

