## Supplementary material for "Quantitative profiling of axonal guidance proteins during the differentiation of human neurospheres"

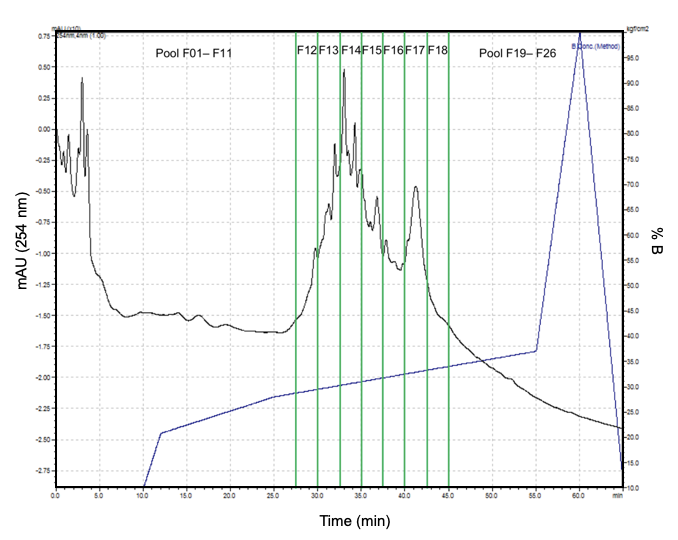

Supplementary figure 1: Offline peptide fractionation in HILIC. Separation was performed in a total analysis of 65 minutes (45 minutes of active gradient). In total, 26 fractions were collected and pooled according to the intensity of the peaks. Fractions F01 to F11 were pooled in a single mix, and fractions F19 to F26 configured another pool, the rest of the fractions were analyzed as single fractions (F12, F13, ...., F18). Thus, a total of 9 fractions (7 individual fractions and 2 pool mix) were further analyzed by LC-MS/MS.

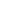
